# Dose-dependent dissociation of pro-cognitive effects of donepezil on attention and cognitive flexibility in rhesus monkeys

**DOI:** 10.1101/2021.08.09.455743

**Authors:** Seyed A. Hassani, Sofia Lendor, Adam Neumann, Kanchan Sinha Roy, Kianoush Banaie Boroujeni, Kari L. Hoffman, Janusz Pawliszyn, Thilo Womelsdorf

## Abstract

**BACKGROUND:** Donepezil exerts pro-cognitive effects by non-selectively enhancing acetylcholine (ACh) across multiple brain systems. The brain systems that mediate pro-cognitive effects of attentional control and cognitive flexibility are the prefrontal cortex and the anterior striatum which have different pharmacokinetic sensitivities to ACh modulation. We speculated that these area-specific ACh profiles lead to distinct optimal dose-ranges for donepezil to enhance the cognitive domains of attention and flexible learning.

**METHODS:** To test for dose-specific effects of donepezil on different cognitive domains we devised a multi-task paradigm for nonhuman primates (NHPs) that assessed attention and cognitive flexibility. NHPs received either vehicle or variable doses of donepezil prior to task performance. We measured donepezil intracerebral and how strong it prevented the breakdown of ACh within prefrontal cortex and anterior striatum using solid-phase-microextraction neurochemistry.

**RESULTS:** The highest administered donepezil dose improved attention and made subjects more robust against distractor interference, but it did not improve flexible learning. In contrast, only a lower dose range of donepezil improved flexible learning and reduced perseveration, but without distractor-dependent attentional improvement. Neurochemical measurements confirmed a dose-dependent increase of extracellular donepezil and decreases in choline within the prefrontal cortex and the striatum.

**CONCLUSIONS:** The donepezil dose for maximally improving attention functions differed from the dose range that enhanced cognitive flexibility despite the availability of the drug in the major brain systems supporting these cognitive functions. Thus, the non-selective acetylcholine esterase inhibitor donepezil inherently trades improvement in the attention domain for improvement in the cognitive flexibility domain at a given dose range.

## INTRODUCTION

The acetylcholinesterase (AChE) inhibitor donepezil (Aricept) is one of few FDA approved cognitive enhancers that aims to address a wide range of cognitive deficits in subjects with mild cognitive impairment or dementia (1–3). Basic research suggests that the cognitive domains that can be enhanced with AChE inhibitors range from selective attention, working memory, response inhibition, learning, and long-term memory (4–6). Consistent with these reports, clinical studies assessing donepezil at one or two doses across larger cohorts of subjects with varying stages of Alzheimer’s disease have found improvements of compound scores of cognitive testing batteries (4,7–10). It is, however, not clear whether the standard doses of donepezil used in clinical studies improve multiple cognitive domains directly, or whether at a particular effective dose its major route of action is to enhance arousal, which then provides an indirect, overall cognitive advantage for attention, working memory, learning and memory processes (6,11). Assessing whether donepezil’s major route of action is the arousal domain, or whether it affects multiple specific cognitive domains simultaneously at a given dose is important for evaluating its therapeutic efficiency and to identify cognitive domains that should be targeted in drug discovery efforts for improved future cognitive enhancers.

One potential limitation of donepezil and other AChE inhibitors is that they increase acetylcholine (ACh) concentrations non-selectively across multiple brain systems. Such a non-selective ACh increase has shortcomings when brain systems are differently sensitive to ACh action so that the same donepezil dose that is optimally affecting one brain system might over- or under-stimulate another brain system. In primates, muscarinic ACh subreceptors relevant for attention and memory functions (12–15), have enhanced densities in prefrontal cortex (PFC) (16), suggesting that PFC may be more sensitive to modulation by AChE inhibitors than posterior brain areas. Moreover, a comparison of transcription factor (CREB) activation of the PFC and the striatum to muscarinic modulation by Xanomeline has reported a 10-fold higher receptor sensitivity of the striatum (17), consistent with other studies reporting significantly higher muscarinic binding potential and higher AChE activity in the striatum than in other cortical regions (18). It is unclear how these differences affect ACh modulation of attention functions that depend on the PFC (19) and on flexible learning functions that are dependent on the striatum (20,21). One consequence of the brain area specific sensitivity to ACh levels could be that a *Best Dose* for enhancing cognitive functions supported by the striatum might not sufficiently stimulate the PFC, and that a *Best Dose* for enhancing PFC functions might overstimulate the striatum.

To test for these possible implications of brain region-specific ACh action, we devised a drug testing paradigm for monkeys that assessed the effects of three different doses of donepezil across different domains of arousal, attention, and cognitive flexibility in a single testing session. We evaluated the attention domain with a visual search task that varied the number and perceptual similarity of distracting objects and quantified the domain of cognitive flexibility with a learning task asking monkeys to flexibly adapt to new feature-reward rules and avoid perseverative responding. This assessment paradigm goes beyond existing nonhuman primate studies of donepezil that so far have found enhanced short-term memory using delayed match-to-sample tasks (4,6,10,15,22–29), enhanced arousal and non-selective speed of processing (15,27), or no consistent effect (18) (**Table S1**). With our multi-domain task design we found that donepezil improves attentional control of interference from distractors at doses that caused an overall slower responding (i.e. reduced speed of processing) and peripheral side effects. In contrast, a lower dose of donepezil caused no clear attentional effect but improved cognitive flexibility. These findings document domain-specific dose-response effects of donepezil for attention and cognitive flexibility.

## METHODS AND MATERIALS

### Nonhuman Primate Testing Protocol

Three adult male rhesus macaques (*Macaca mulatta*; ∼8-15 kg, 6-9 years old) were used for this experiment. They were separately given access to a cage-mounted Kiosk Station that provided a touchscreen interface inside the animal’s housing unit to perform cognitive tasks (**Figure 1A**) (30). Monkeys were cognitively assessed at the same time of day for ∼20ml/kg fluid reward. The behavioral tasks, reward delivery, and the registering of behavioral responses were controlled by the Unified Suite for Experiments (USE) (31). The task protocols, matlab analysis procedures and the open-sourced USE software are available at http://accl.psy.vanderbilt.edu/resources/analysis-tools/unifiedsuiteforexperiments/.

**Figure 1.**
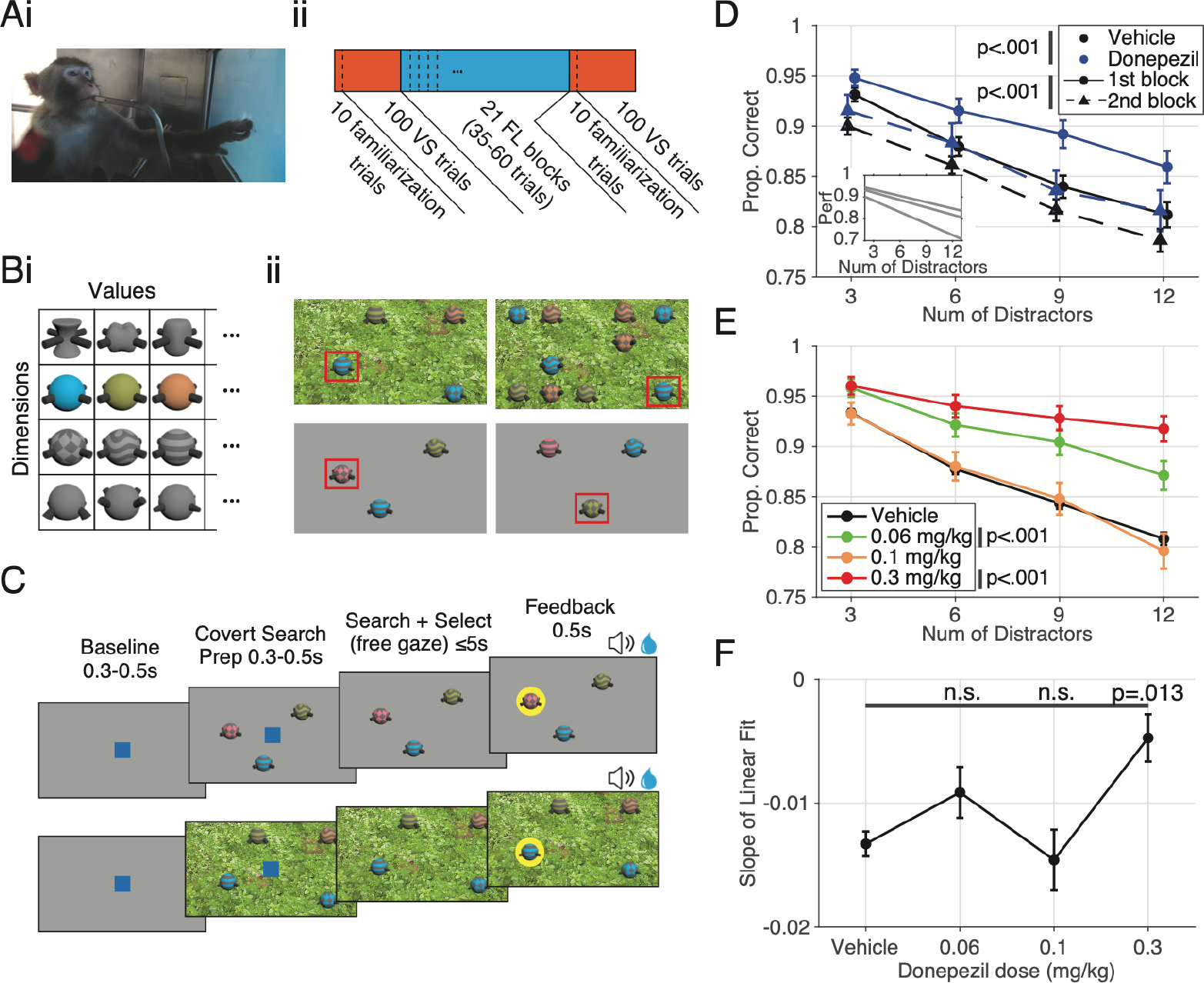
Task design, meta-structure and visual search performance as a function of distractor number. (**A**) **i**. Picture of one of the subjects working in the custom-built kiosk, interacting with the touchscreen and receiving fluid reward. **ii**. The meta structure of the Multi-task. Each experimental session consists of 3 super-blocks of VS, FL and VS respectively. Each VS block is preceded by 10 familiarization trials which is identical to a VS trial but without any distractors. Each VS block contains trials with 3/6/9/12 distractors randomly selected and counterbalanced over the block. In contrast, each FL block will contain 0/1 irrelevant feature dimensions in addition to the relevant feature dimension (the dimension with the rewarded feature value) counterbalanced over the session. (**B**) **i**. From the grand pool of quaddles which includes four feature dimensions and a variable number of feature values (9 shapes, 9 patterns, 8 colors, and 11 arms), three feature values from three feature dimensions are chosen. This 3×3 pool is then counterbalanced for dimension presentation and feature reward association and is utilized for 2 weeks of data collection where all presented quaddles are selected from this 3×3 pool. **ii**. Example trials. Two example VS trials (top) within the same block with 3 distractors (left) and 9 distractors (right). Each VS block will contain one of 5 backgrounds, with the VS blocks in the same day never having the same background. All distractors and target objects in VS blocks are three dimensional objects and distractors may be duplicated in each trial. Quaddles are spatially randomly presented at the intersections of a 5×4 virtual grid pattern on screen. The red box highlights the rewarded target object, which is invariable within the VS block, in these examples. Two example FL trials (bottom) within the same block containing 2D quaddles (1 distracting dimension plus the relevant dimension). The rewarded feature value in this block is the checkered pattern independent of what color feature value it is paired with. Quaddles may be presented in 8 possible locations in a circle each being 17 degrees of visual radius away from the center of the screen. The red box signifies the rewarded target object, which is a variable combination of the rewarded feature value (the checkered pattern in this example) with a random feature value of the distractor dimension (color in this example). (**C**) The trial structure for both the FL (top) and VS (bottom) blocks of the task are very similar. A trial is initiated by a 0.3-0.5s touch and hold of a blue square (3° visual radius wide) after which the blue square disappears for 0.3-0.5s before task objects, which are 2.5° visual radius wide, are presented on screen. Once the task objects are on screen, the subject is given 5s to visually explore and select an object via a 0.2s touch and hold. A failure to make a choice within the allotted 5s results in an aborted trial and did not count towards the trial count. Brief auditory feedback and visual feedback (a halo around the selected object) are provided upon object selection, with any earned fluid reward being provided 0.2s following object selection and lasting 0.5s along with the visual feedback. Non-rewarded trials had a different auditory tone and a light blue halo around the selected, non-rewarded object. Rewarded objects had a higher pitch auditory tone, a light yellow halo around the selected rewarded object and had an accompanying fluid (water) reward. (**D**) Average VS performance by distractor number for vehicle and all donepezil doses combined, both separated by the first vs second VS block. VS performance was significantly different for block number (F(1,1722) = 22.19, p < .001) as well as condition (F(1,1722) = 19.0, p < .001). The inlet shows individual monkey average VS performance linear fits. **(E**) Average VS performance by distractor number between vehicle and 0.06, 0.1, and 0.3 mg/kg donepezil doses for the first VS block (p < .001). Both the 0.06 and 0.3 mg/kg doses were significantly different from vehicle (p < .001). Error bars here reflect standard deviation in this panel. (**F**) The set size effect of VS performance by distractor number for each condition. The 0.3 mg/kg dose set size effect was significantly shallower from the vehicle set size effect (H(3) = 11,46, p = .010; Tukey’s, p = .013).

### Drugs and Procedures

Donepezil-hydrochloride was purchased from Sigma-Aldrich (catalog number D6821; St. Louis, MO, USA). We tested three doses of donepezil: 0.06, 0.1 and 0.3 mg/kg to operate within the dosing range of previous studies reporting pro-cognitive effects (**Table S1**). At this IM range, plasma concentrations of donepezil have been shown to be roughly the same when dosing with ∼10x the concentration via PO (15). All drug doses were administered in a double-blind manner. Animals received saline as vehicle control, or a dose of donepezil IM injection 30 minutes prior to starting task performance taking into account its expected 1h half-life (32). Drug side effects were assessed 15 min following drug administration and after completion of the behavioral performance with a modified Irwin Scale (33–36) for rating autonomic nervous system functioning (salivation, etc.) and somato-motor system functioning (posture, unrest, etc.). Monkeys’ behavioral status was video-monitored throughout task performance (**Figure 1A**).

### Behavioral Paradigms

Monkeys performed a visual search (VS) task to measure attentional performance metrics and a feature-reward learning (FL) task to measure cognitive flexibility metrics in each experimental session. Each performance day was made up of an initial set of 100 trials of VS, a set of 21 learning blocks with 35-60 trials each of the FL task, and a final set of 100 trials of the VS task (**Figure 1Aii**). Details of both tasks are provided in the **Supplement**. In brief, the VS task required monkeys to find and touch a target object among 3, 6, 9, or 12 distracting objects in order to receive fluid reward (**Figure 1B**). The target was the object that was shown in 10 initial trials without distractors. Targets and distractors were multidimensional, 3D rendered Quaddle objects (31) that shared few or many features of different features dimensions (colors, shapes, arms, body patterns), which rendered search easier when there was no or few similarities among features of targets and distractors, or more difficult if the target-distractor (T-D) similarity was high (**Figure 2A**). The FL-task required monkeys to learn through trial-and-error which object feature is rewarded in a given block of ∼35-60 trials (**Figure 1C**). In each trial of the block three objects were shown that varied either in features of one feature dimension (e.g. having different colors or different body shapes), or that varied in features of two feature dimensions (e.g. having different colors and different body shapes). Choosing the object with the correct feature was rewarded with a probability of 0.8. Blocks where only 1 feature dimension varied (*1D blocks*) were easier as there was lower attentional load than in blocks with 2 varying feature dimensions (*2D blocks*).

**Figure 2.**
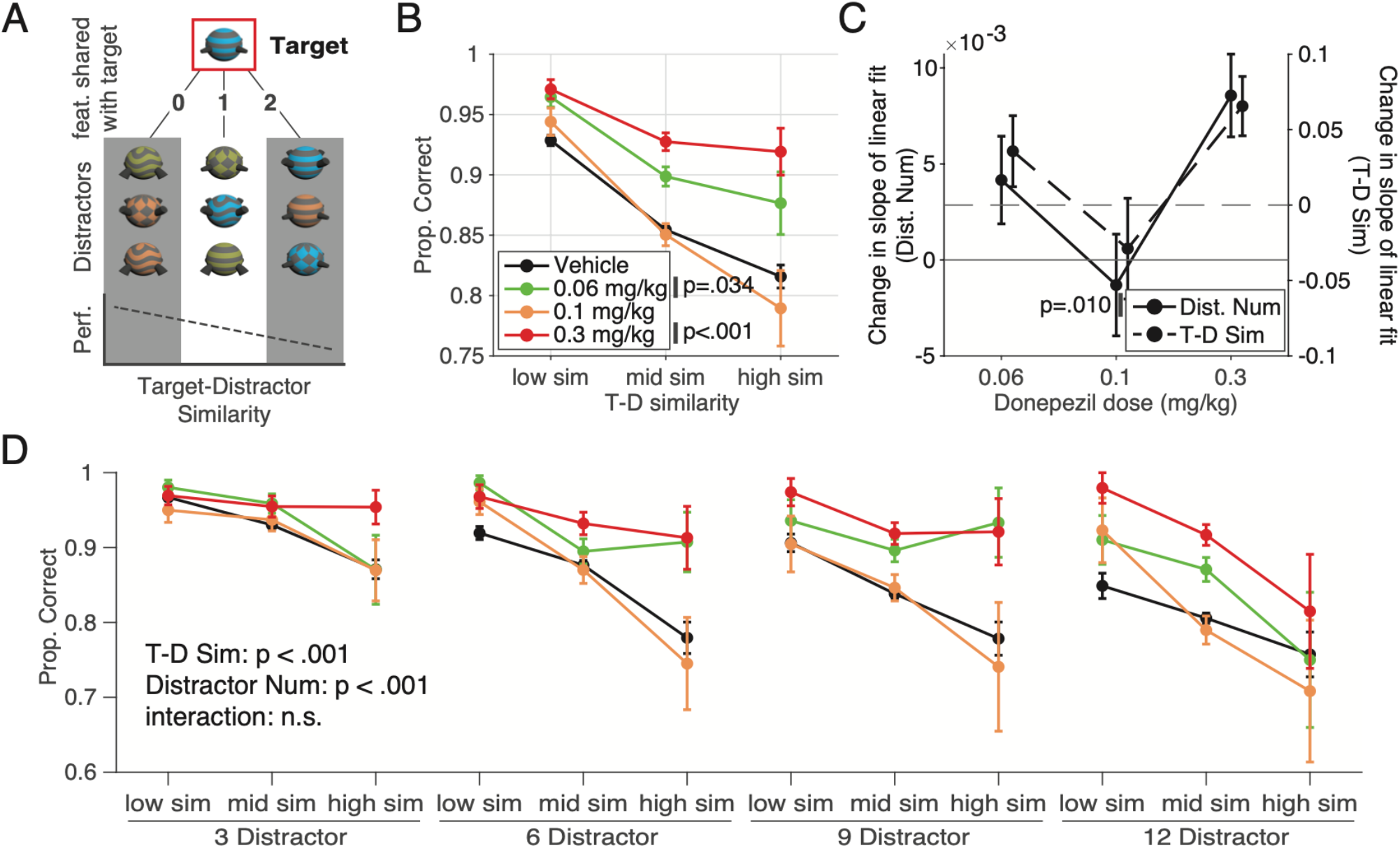
Visual search task performance and change in difficulty through increasing distractor numbers and target-distractor similarity. (**A**) A visual description of the target-distractor similarity measure in the VS task. For an example target in the red square, 3 example distractors are presented with 0, 1 and 2 features in common respectively from left to right. The cartoon plot below shows the impact of the average target-distractor similarity of an individual trial on performance. (**B**) Similar to **Figure 1D**, but here we plot average VS performance by T-D similarity. There was a significant effect of T-D similarity on performance (F(2,627) = 16.17, p < .001) as well as condition with both the 0.06 and 0.3 mg/kg donepezil doses being significantly different from vehicle (F(3,267) = 7.75, p < .001; Tukey’s p = .034 and p < .001 respectively). (**C**) The change in the slope of VS performance with 0.06, 0.1 and 0.3 mg/kg donepezil relative to vehicle. The change in slope by distractor number is plotted on the left y axis (same data as **Figure 1F**) (H(3) = 11.46, p = .010) while the change in slope by T-D similarity is plotted on the right y-axis (H(3) = 2.8, n.s.). (**D**) A visualization of the combined effect of distractor number and T-D similarity on performance. From left to right, each cluster of lines represents increasing distractor numbers while data within each line represents low, medium and high T-D similarity from left to right respectivel. Both distractor number (F(3,16688) = 50.25, p < .001) and T-D similarity (F(2,16688) = 55.24, p < .001) impact VS performance with no significant interaction (F(6,16688) = 1.16, n.s.).

### Neurochemical Confirmation of Drug Effect

To evaluate the levels of donepezil in brain structures that are necessary for successful attention and learning performance, we measured choline and donepezil concentrations in the prefrontal cortex and the anterior striatum (caudate nucleus) 15 min after administering a low and high dose of donepezil (0.06 and 0.3 mg/kg, IM) in a separate experiment. Measures of donepezil were made at the time when we observed dose-limiting side effects at the 0.3 mg/kg dose and the two tested doses were accompanied by pro-cognitive effects in our task (see results). We used microprobes that sampled the local neurochemical milieu with the principles of solid phase micro-extraction (SPME) (for details see **Supplement**) (37). SPME probes sampled the level of donepezil and the ACh metabolite (choline) via diffusion at a consistent rate until an equilibrium was reached with the extracellular concentrations. The neurochemical concentrations were quantified with liquid chromatography and mass spectrometry as described in detail in (37). The detailed procedures used here are described in (38).

### Statistical Analysis

Data were analyzed with standard nonparametric and parametric tests as described in the **Supplement**.

## Results

Each monkey was assessed in 38 sessions total including 17 vehicle days and 7 days with each dose (0.06, 0.1 and 0.3 mg/kg). Drug side effects were noted following IM injections of the 0.3 mg/kg dose in the first 30 min post injection as changes in posture, sedation, vasoconstriction and paleness of skin, but no adverse effects persisted beyond 30 min. (**Table S2**). First, we confirmed that monkeys performed the visual search (VS) task at high 84.4% (± 0.54) accuracy (monkeys Ig: 85.2% ±0.81; Wo: 88.3% ±0.94; Si: 79.8% ±0.97) and showed the expected set-size effect evident in decreased accuracy and slower reaction times with increasing numbers of distractors (**Figure 1D, Figure S1** and **S2, Supplemental**). When targets were more similar to distractors (high T-D similarity) VS performance decreased from 92.9% (±0.4) to 85.5% (±0.3) and 81.6% (±1.0) for low, medium and high T-D similarity, respectively (H(2) = 169.48, p < .001) (**Figure 2B**). In the feature learning (FL) task, the monkeys reached learning criterion faster in the easier 1D (low distractor load) condition (avg. trials to ≥80% criterion: 12.5 ± 0.2 SE), than in the 2D (high distractor load) condition (avg. trials to ≥80% criterion: 15.6±0.2) (**Figure 3A, Supplemental**).

**Figure 3.**
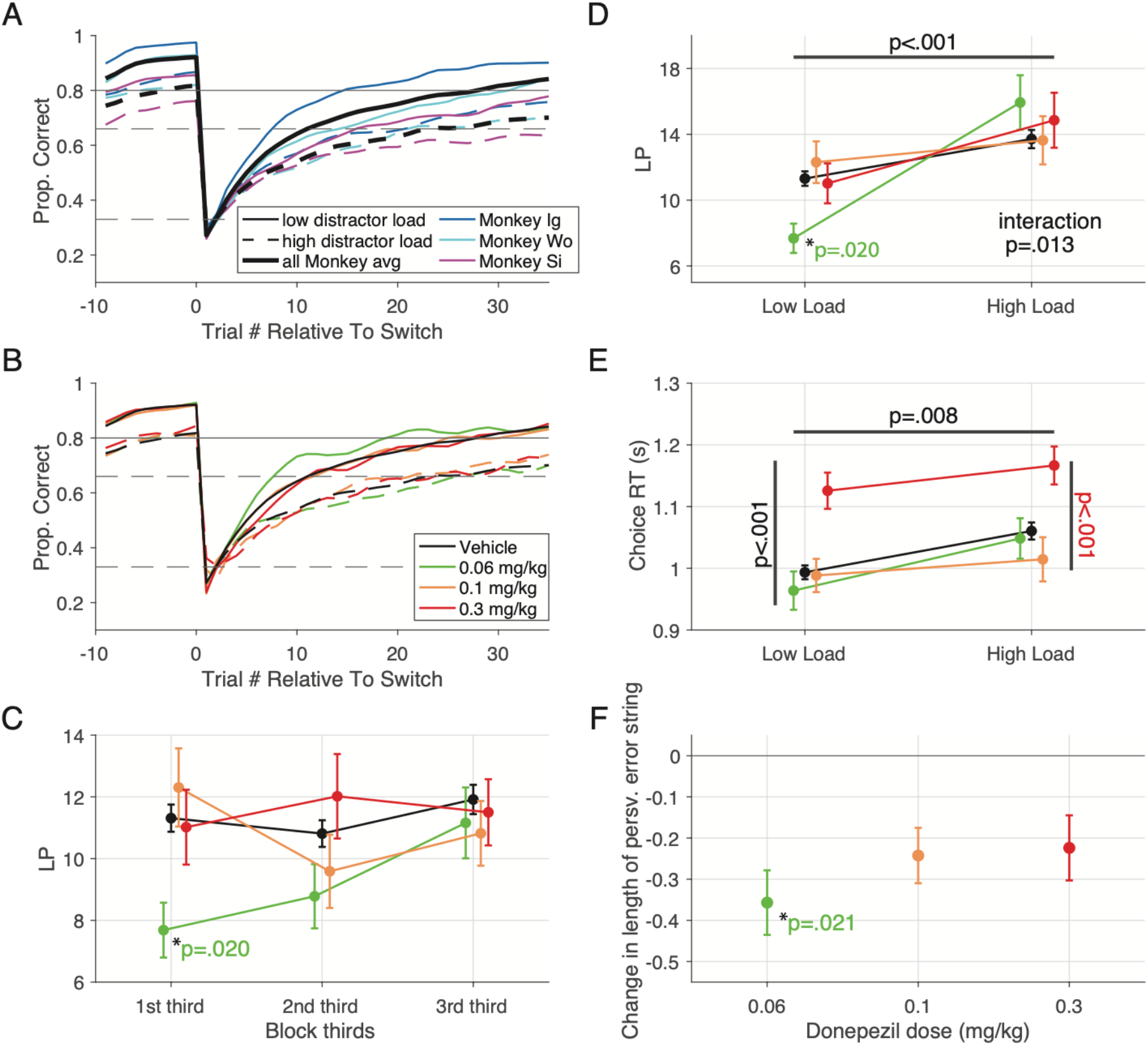
Feature learning task learning curves and performance. (**A**) Average learning curves of each monkey and all monkeys combined for both low and high distractor load conditions. In all instances, monkeys learned faster and with higher plateau performance in low distractor load blocks relative to high distractor load blocks. (**B**) All monkey average learning curves for vehicle, 0.06, 0.1 and 0.3 mg/kg donepezil doses for both low and high distractor load conditions. (**C**) Temporal progression of learning speed (LP) for vehicle, 0.06, 0.1 and 0.3 mg/kg donepezil doses for the low distractor load condition only. At the 0.06 dose, donepezil allows for faster learning in the low attentional load blocks (F(3,602) = 3.3, p = .020). Similar to the VS task, donepezil’s enhancement is only visible early on and relatively close to its i.m. administration. (**D**) Average learning speed of vehicle and donepezil doses for low and high distractor load blocks across sessions reveals an interaction between drug condition and distractor load (F(3,1052) = 3.59, p = .013). (**E**) The same as D but for choice RTs instead of learning speed. The 0.3 mg/kg donepezil dose slows choice reaction times in both low and high distractor load blocks (Condition F(3,1052) = 12.3, p < .001; Tukey’s, p < .001). (**F**) Change in the length of perseverative errors from vehicle, where feature values in the distracting dimension were the target of the perseverations. Error bars reflect SEMean for inter-monkey variability. Donepezil at the 0.06 mg/kg dose significantly reduces perseveration length in the distracting dimension (p = .021); other donepezil doses trends towards this as well.

### Dose-dependent improvement of visual search accuracy and slowing of choice reaction times

Donepezil significantly improved accuracy of the visual search task (F(1,1722) = 18.95, p < .001)(**Figure 1D**), but on average slowed search reaction times (F(1,1722) = 4.83, p = .028)(**Figure S1B**). The slower choice reaction times were evident already to the single target object in the 10 target familiarization trials (**Figure S1A**). These main behavioral drug effects were evident prominently in the first visual search block (**Figure 1D, Figure S1A**). We therefore focused our further analysis on the first search block.

The improved accuracy of visual search was dose-dependent. The 0.1 mg/kg dose enhanced performance by 2.5% ±1.0, 4.4% ±1.3, 6.1% ±1.4 and 6.3% ±1.6 (mean ±SD) for 3/6/9/12 distractor trials, respectively (*X*^2^(1, N_1_ = 16700, N_2_ = 2100) = 35.5, p < .001). The 0.3 mg/kg dose enhanced performance by 2.7% ±1.0, 6.3% ±1.2, 8.5% ±1.3 and 11.0% ±1.4 (mean ±SD) for 3/6/9/12 distractor trials respectively (*X*^2^(1, N_1_ = 16700, N_2_ = 1900) = 75.9, p < .001) (**Figure 1E**). Thus, we found larger improvement the more distractors interfered with the target search. We confirmed this by fitting a regression line across performance at different number of distractors, which revealed overall significantly shallower slopes with donepezil (slopes: -0.013 ±0.001, - 0.009 ±0.002, -0.015 ±0.003 and -0.005 ±0.002 for vehicle, 0.06, 0.1, and 0.3 mg/kg of donepezil respectively (H(3) = 11.46, p = .01)). Pairwise comparison showed that the 0.3 mg/kg drug dose and the vehicle condition showed significantly different slopes (Tukey’s, p = .013)(**Figure 1F**).

In contrast to improving visual search accuracy, donepezil slowed down reaction times across all distractor conditions at the 0.3mg/kg dose relative to vehicle by on average 100 ms ±40, 238 ms ±79, 208 ms ±99, 264 ms ±102 (mean ± SD) for 3/6/9/12 distractors respectively (p = .023, Bonferroni correction)(**Figure S1C**). The slope of the regression over different number of distractors did not differ between 0.3 mg/kg dose and vehicle which shows the reaction time effect is a non-selective effect that is independent of distractors (regression slope on RTs: 0.061 ±0.002, 0.065 ±0.007, 0.067 ±0.007 and 0.076 ±0.009 (H(3) = 3.37, n.s.) for vehicle, 0.06, 0.1, 0.3 mg/kg of donepezil respectively (**Figure S1D**).

Across sessions visual search accuracy was correlated with reaction times only for the vehicle (Pearson, ρ: -0.30, p < .001) and 0.1 mg/kg donepezil dose condition (Pearson, ρ: -0.46, p = .034), but not for the 0.06<u>and 0.3 mg/kg</u> dose conditions in which monkeys showed improved accuracy, which suggests the accuracy improvement is independent from a slowing of reaction speed (**Figure S2A-B**).

We next tested whether improved control of interference from increasing number of distractor objects was likewise evident when increasing the similarity of distractor and target features (**Figure 2A**). First, we confirmed that higher target-distractor similarity overall reduced performance (F(2,672) = 16.17, p < .001, **Supplemental**). Donepezil significantly counteracted this similarity effect and improved performance at the 0.06 and 0.3 mg/kg doses (F(3,672) = 7.75, p < .001, Tukey’s, p = .034 and p < .001 respectively). This finding shows that the beneficial effect of donepezil significantly increased when there was higher demand to control perceptual interference from distracting objects (**Figure 2B**). This was also evident as a statistical trend of a shallower regression slope at 0.06 and 0.3 mg/kg doses of donepezil, which indicates less interference from distracting features when they were similar to the target (**Figure 2C**) (H(3) = 2.79, n.s.; slope changes relative to vehicle for 0.06, 0.1 and 0.3 mg/kg doses were: +0.0357 ±0.0236, -0.0289 ±0.0334, -0.0656 ±0.0197). The improved search performance with donepezil for visual search with higher target-distractor similarity and with a higher number of distractors was evident in significant main effects, but there was no interaction, suggesting they improved performance independently of each other (F(2, 16688) = 55.24, p < .001; F(3,16688) = 50.25, p < .001; F(6,16688) = 1.16, n.s. respectively)(**Figure 2D**). This independence was also suggested by the absence of a correlation of the target-distractor similarity effect and the number-of-distractor effect (Pearson, n.s.) (**Figure S3**).

### Dose-dependent improvement of flexible learning performance

Donepezil also improved feature learning performance, but only at the 0.06 mg/kg dose (**Figure 3B**) and most pronounced for the first third of the behavioral session (F(3,602) = 3.3, p = .020; **Figure 3C**). We therefore focused further analysis on the first third of the learning blocks, which revealed that the learning improvement at the 0.06 mg/kg dose was significant for the low distractor load condition (significant interaction effect of drug condition and distractor load (Condition x Distractor Load F(3, 1052) = 3.59, p = .013); and for vehicle, 0.06, 0.1 and 0.3 mg/kg donepezil doses the trials to criterion were 11.3 ±0.4, 7.7 ±0.9, 12.3 ±1.3 and 11.0 ±1.2 trials long with the 0.06 mg/kg dose and vehicle being significantly different (p = .020, Bonferroni correction)(**Figure 3D**). There was no change in learning speed with other doses at low or high distractor load.

Beyond learning speed, we found overall slower choice reaction times at the 0.3 mg/kg donepezil dose (**Figure 3E**) (main effect of drug condition: (F(3,1052) = 12.29, p < .001). While reaction times were overall slower at high distractor load (F(1,1052) = 7.18, p = .008) there was no interaction with drug dose (F(3,1052) = 0.26, n.s.). After visually inspecting the results we separately tested the 0.3 mg/kg dose of donepezil and found it led to significantly slower choice reaction time than vehicle (Tukey’s, p < .001)(**Figure 3E**). The changes in choice reaction times did not correlate with changes in learning performance (number of trials to criterion) at any drug condition, indicating they were independently modulated (Pearson, all n.s.)(**Figure S2D**).

We predicted that the faster learning at the 0.06 mg/kg donepezil dose could be due to a more efficient exploration of objects during learning, which would be reflected in reduced perseverative choices of unrewarded objects. Overall, perseverative errors (defined as consecutive unrewarded choices to objects with the same feature dimension) made up 20% of all errors. As expected, we found significantly shorter sequences of perseveration of choosing objects within distractor feature dimensions at the 0.06 mg/kg dose of donepezil (**Figure 3F**). For 0.06, 0.1 and 0.3 mg/kg doses the average length of perseverations in the distractor dimension was: 2.1 ±0.1, 1.8 ±0, 1.9 ±0.1 and 1.9 ±0.1 trials with the difference between vehicle and the 0.06 dose being significant (p = .021). Perseverative choices in the target feature dimension were not different between conditions (for 0.06, 0.1 and 0.3 mg/kg donepezil doses the avg. perseveration length in the target dimension was: 1.7 ±0, 1.7 ±0, 1.6 ±0, and 1.7 ±0 trials (n.s.).

### Dissociation of attention and learning improvements, but slowing is correlated

The effects of donepezil on feature learning and visual search might be related, but we found that learning speed and search accuracy was not correlated at those doses at which the drug improved learning and search (0.06 mg/kg dose) or improved only visual search (0.3 mg/kg dose) (Pearson, all n.s.). A significant correlation was found only for the 0.1 mg/kg dose (Pearson, ρ: -0.54; p = .012) (**Figure 4A**). Learning at low or high distractor load and the set size (slope) effects in the visual search task was uncorrelated (Pearson, all n.s.). However, at the 0.3 mg/kg donepezil dose we found that the target-distractor similarity effect (i.e. the search slope change) in the visual search task was positively correlated with the difference of the learning speed at high versus low distractor load in the learning task (Pearson, ρ: 0.60; p = .008). This effect signifies that better attentional search of a target among similar distractors is associated with poorer flexible learning of new targets when there are multiple object features to search through (high distractor load).

**Figure 4.**
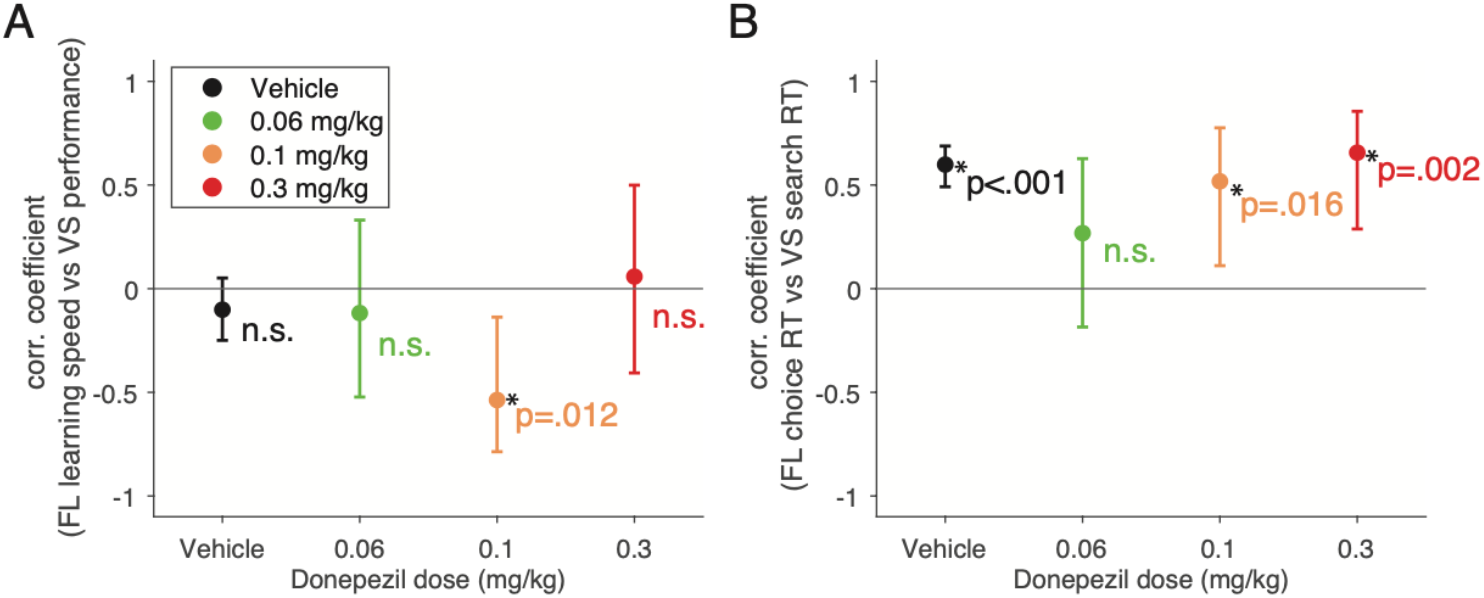
The relationship between the visual search task and the feature learning task. (**A**). Correlation coefficients between FL learning speed (LP) and VS performance for vehicle, 0.06, 0.1 and 0.3 mg/kg donepezil doses. Only the 0.1 mg/kg donepezil dose had a significant correlation between FL and VS task performance (Pearson, ρ: -0.54; p = 0.012). No doses showed a significant change in correlation from vehicle. (**B**) Same as figure A but for FL choice RTs and VS search RTs. Although vehicle, 0.1 and 0.3 mg/kg donepezil doses had a significant correlation between choice and search RTs, we found no significant change in correlation relative to vehicle.

In contrast to accuracy, choice reaction times in the learning task and visual search were significantly correlated for the 0.1 mg/kg donepezil dose (Pearson, ρ: 0.52; p = .016), the 0.3 mg/kg dose (Pearson, ρ: 0.66; p = .002), and the vehicle control condition (Pearson, ρ: 0.60; p < .001)(**Figure 4B**).

### Determination of extracellular donepezil and choline levels in the prefrontal cortex and anterior striatum

Visual search and flexible learning are realized by partly independent brain systems, including the PFC and anterior striatum (39). To determine whether extracellular levels of donepezil were increased to a similar magnitude in the PFC and anterior striatum, we measured its concentration after administering doses of either 0.06 and 0.3 mg/kg donepezil IM in the PFC, assumed to be necessary for efficient interference control during visual search (19), and in the head of the caudate nucleus which is necessary for flexible learning of object values (20,21). We used a recently developed microprobe that samples chemicals in neural tissue based on the principles of solid-phase microextraction (SPME) (37,38). We found that donepezil was available in both brain areas and its extracellular concentration more than doubled after injecting 0.3 mg/kg than 0.06 mg/kg in both areas (F(1,16) = 9.69, p = .007), with no significant difference between PFC and caudate (F(1,16) = 1.44, n.s.)(**Figure 5A**). Donepezil should cause a depletion of the ACh metabolite choline (40). Using HPLC/MS analysis of the SPME samples we found in the PFC that 0.06 and 0.3 mg/kg donepezil reduced choline concentrations by 74.2% ±14.9 (p = .005) and 85.7% ± 26.9 (p = .007) of their baseline concentrations, and in the caudate, it reduced choline by 68.4% ±13.8 (p = .022) and 81.0% ±12.9 (p = .009) of respective baseline concentrations (**Figure 5B**). The 11.5% and 12.6% stronger reduction choline at the 0.3 versus 0.06 mg/kg dose in PFC or caudate was not significant (n.s.).

**Figure 5.**
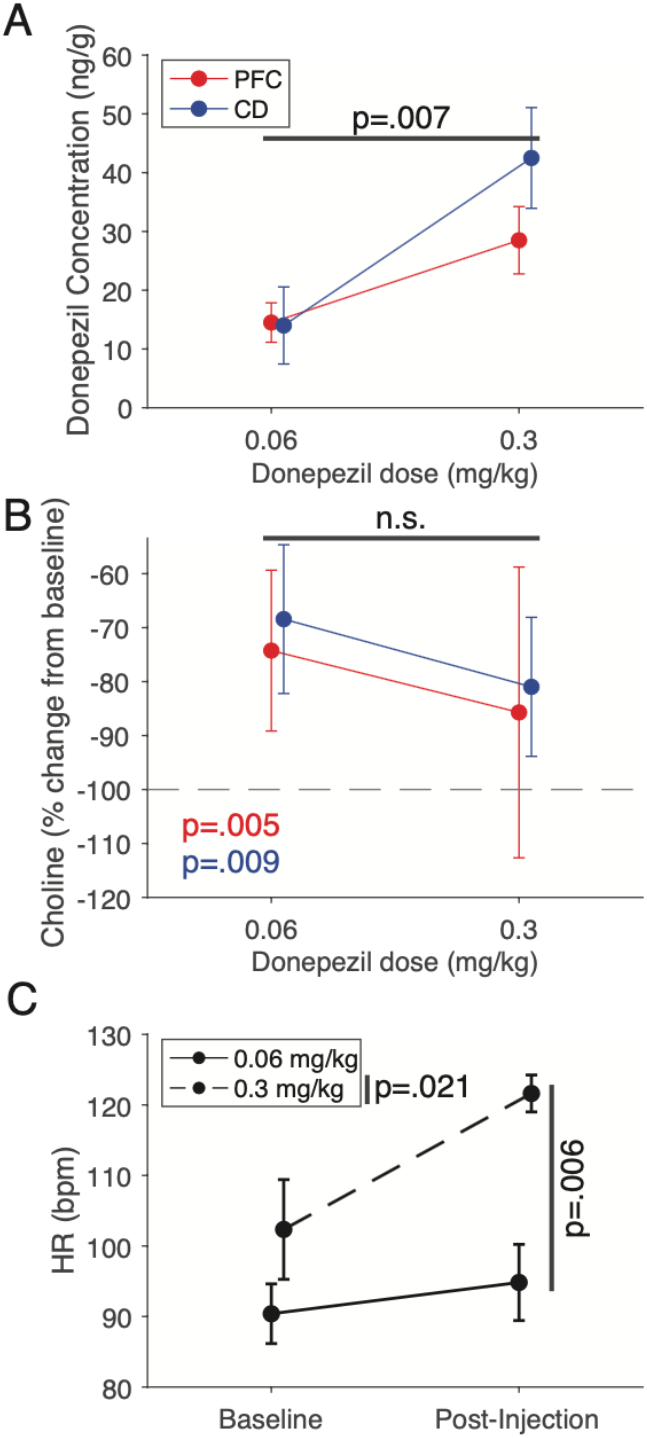
In-vivo extracellular measurements of choline, donepezil as well as donepezil’s effect on heart rate. (**A**) Quantified concentration of extracellular unbound donepezil with 0.06 and 0.3 mg/kg donepezil administration in the PFC and CD. We are able to reliably detect higher donepezil concentrations with 0.3 mg/kg dosing relative to 0.06 mg/kg dosing (Condition F(1,16) = 9.69, p = .007) with SPME. We also see a trend towards higher detectable donepezil in the caudate relative to the PFC at the 0.3 mg/kg dose tested, however, we found neither significant group or interaction effects. (**B**) We used choline concentrations as a metric for donepezil bio-activity as it de-activates AChE and prevents acetylcholine’s degradation into choline. We extracted average session-wise change in choline from baseline with 0.06 and 0.3 mg/kg donepezil doses within the PFC and CD. Although we find significant decreases in choline by up to >80% of baseline concentrations, we found no significant effect of dosing in either the PFC or CD. (**C**) The heart rate of our fourth monkey was monitored during the neurochemical experiments. This revealed a sharp and transient increase in HR post administration of donepezil at 0.3 mg/kg dose (**Supplemental**) which lead to a higher average bpm. We found that we can significantly distinguish 0.06 and 0.3 donepezil administration via HR (p = .006).

To obtain an independent physiological marker of dose-dependent effects we quantified during actual task performance how donepezil changed the heart rate (HR) before versus after drug administration (**Supplemental**). HR showed a transient peak ∼20 min after donepezil injection relative to baseline, which was significant for the 0.3 mg/kg dose (pre-injection 102.3 ±7.1 to post-injection 121.6 ±2.6; p = .021), but not for the 0.06 mg/kg dose (pre-injection: 90.3 ±4.2 to post-injection: 94.8 ±5.4; n.s.). The 0.3 mg/kg dose caused a significantly higher HR peak than the 0.06 mg/kg dose (p = .006) (**Figure 5C**).

## Discussion

Here, we dissociated donepezil’s improvement of attentional control of interference during visual search performance from improvements of cognitive flexibility during feature reward learning. At the highest dose tested donepezil reduced interference during visual search particularly when there were many distractors and high similarity of distractors to the target, while concomitantly slowing down overall reaction times and inducing temporary peripheral side effects. In contrast, at the lowest dose donepezil did not affect target detection times during visual search, but improved adapting to new feature-rules and reduced perseverative responding. These findings document a dose-dependent dissociation of the best dose of donepezil for improving attention and for improving cognitive flexibility.

### Different donepezil dose-ranges for improving interference control and flexible learning

Using a behavioral assessment paradigm with two tasks allowed us to discern differences of the donepezil dose that maximally improved interference control (in the visual search task) versus the dose that maximally improved flexible learning (in the reward learning task). In both tasks, donepezil modulated performance early within the session (first of two VS blocks and first third of FL blocks) consistent with its short half-life and rapid time to peak concentration with IM delivery (15,32) and therefore our results focused on this time window. At the 0.06 mg/kg dose donepezil facilitated flexible learning of a new feature reward rule and reduced the length of perseverative errors (**Figure 3C,F**). These behavioral effects can be interpreted as improvements of cognitive flexibility of the monkeys in adjusting to changing task demands. At the same 0.06 mg/kg dose visual search response times were unaffected (**Figure S1**) and visual search accuracy was overall improved but independent of the number of distractors, i.e. independent of the degree of interference (**Figure 1E,F**). In contrast, at the higher donepezil doses flexible learning behavior was indistinguishable from the no-drug vehicle control condition showing that improving flexibility required donepezil at a lower dose.

This conclusion is opposite to the drug effects on visual search performance, which was maximally improved at the 0.3 mg/kg dose. At this dose, subjects not only showed improved resistance to interference when there were more distracting objects (**Figure 1E,F**), but also improved resistance to distracting objects that were visually similar to the searched-for target (**Figure 2B-D**). These findings document that donepezil enhances the robustness to distraction (41,42), which critically extends insights from existing primate studies with donepezil that mostly used simpler tasks to infer pro-cognitive effects on working memory or arousal (*see* **Table S1**). The process of attentional control of interference goes beyond a short-term memory effect measured with delayed match to sample tasks. In the visual search paradigm we used, short-term memory of the target object is already necessary for performing the easier trials with 3 or 6 distractors, while an attention specific effect can be inferred when there is greater improvement in performance with increased attentional demands in trials with 9 or 12 distractors. Thus, our study provides strong evidence that donepezil can cause specific attentional improvement at relatively higher doses. This finding supports a prominent neuro-genetic model of cholinergic modulation of attention (43) that has received recently functional support in studies reporting enhanced distractor suppression in nonhuman primates with nicotine receptor specific ACh modulation (44–46), and improved suppression of perceptually distracting flankers in human subjects tested with a single dose (47).

### Non-selective slowing of response times and dose-limiting side effects

We found that 0.3 mg/kg donepezil overall slowed response times of the monkeys during visual search independent of distractor number or target-distractor similarity (**Figure S1A,C**), and during feature-reward learning independent of distractor load (**Figure 3E**). The slowing of reaction times was independent of overall accuracy levels (**Figure 4A**), which shows it did not reflect trading off speed for accuracy. The observed slowing occurred at a dose that improved attention and was unexpected, because prior studies using the delayed match to sample task did not report changes in reaction times in monkeys (23,25), or reported normalized reaction times in studies using donepezil to recover from scopolamine induced deficits (15,26) (**Table S1**). Our findings therefore indicate that 0.3 mg/kg of donepezil already induced cholinergic side effects while still improving cognitive processes. This interpretation is supported by our observation of arousal deficits at the 0.3 mg/kg dose that became apparent in vasoconstriction, changes in posture, visible sedation and paleness (**Table S2**). These side effects were strongest within 30 min. after administration of the drug. Although these side effects did not prevent animals from starting and completing the tasks, they limited the dose range we could test. Such dose-limiting side effects are a well known limitation of donepezil and other AChE inhibitors where therapeutically effective doses cause in a subset of patients gastrointestinal issues such as nausea, diarrhea, and arousal deficits (10,48–50). Our finding adds to this literature that arousal deficits are occurring at a dose range that causes apparent improvements in attentional control of interference while lower doses that were void of side effects failed to improve attention. These observations might have clinical implications as they predict that lower doses of donepezil might not cause improved attention, but primarily improve cognitive flexibility.

Our finding of dose-limiting side effects and reductions in arousal or speed-of-processing emphasizes the importance of developing drugs that avoid nonselective overstimulation of intrinsic cholinergic neurotransmission. Strong candidate compounds include positive allosteric modulators (PAMs) for nicotinic subreceptors (51) and for M_1_ and M_4_ muscarinic receptor (13,14,52,53). The subtype-specific muscarinic PAMs do not target the orthosteric binding site of acetylcholine (ACh) that is highly conserved across all five mAChR subtypes, but rather, they act at more topographically distinct allosteric sites. In addition, PAMs have no intrinsic activity at their respective muscarinic receptor subtype, but act to boost normal cholinergic signaling thereby conserving the spatial and temporal endogenous ACh signaling and avoiding overstimulation of peripheral ACh receptors and subsequent adverse side effects (54–56). Our study thus provides an important benchmark for the development of new drugs that aim to enhance multiple cognitive domains while minimizing side effects.

### Quantifying extracellular levels of donepezil and choline in prefrontal cortex and striatum

We confirmed the presence of extracellular donepezil in the PFC and the anterior striatum at the doses tested (**Figure 5A**) and that it prevented ACh metabolism as evident in 68-86% reduced choline levels (**Figure 5B**). To our knowledge this is the first quantification of donepezil’s action on the breakdown of ACh in two major brain regions in the primate. The observed reduction of choline is higher than reductions of AChE activity (of ∼25-70%) reported with positron emission tomography or in brain homogenate (57,58). Previous studies suggest that evaluating blood plasma levels or cerebrospinal concentrations may not predict how effectively drugs acting on AChE influence behavioral outcomes (59). One likely reason is that intracerebral concentrations can be multifold higher than extracerebral concentration levels (57,60) and do not reflect the actual bioactive concentration available in target neural circuits. By confirming that donepezil prevented ACh breakdown in the PFC and striatum, we thus established a direct link of behavioral outcomes and local drug action in two brain structures whose activity causally contributes to attention and learning. The lateral PFC is causally necessary for attentional control of distractor interference (61) with ACh depletion of the PFC impairing attention, but not learning (62). In contrast, flexible learning and overcoming perseverative response tendencies closely depend on the anterior striatum (63). Both structures closely interact during attention demanding learning processes (64), but can be dissociated neurochemically (39). Our findings showed that donepezil has a similar effect on ACh breakdown in both areas, which suggests that differences in behavioral outcomes at a given donepezil dose are likely due to differences in the sensitivity of these areas to ACh action. Indeed, prior studies suggest that the striatum has a particularly high muscarinic binding potential (18) and respond (tenfold) stronger to muscarinic ACh receptor activation compared with the PFC (17). We speculate that these brain area specific neuromodulatory profiles underly the observed dose specific improvements of cognitive flexibility and attentional control of interference.

The neurochemical measurements of donepezil in PFC and striatum were achieved with a recently developed microprobe that samples neurochemicals through principles of solid phase microextraction (SPME) (37,38,65–67), and so far was used for testing the consequence of drugs only in rodents (66,68,69). We believe that leveraging this technique in primate drug studies will be important for clarifying whether systemically administered drugs reach the desired target brain systems in which they are supposed to exert their pro-cognitive effects.

In our study, confirming donepezil’s action in PFC and striatum critically constrains the interpretation of the behavioral results, suggesting that different behavioral outcome profiles are not due an uneven drug availability. Rather, the different ‘*Best Doses*’ for visual search and flexible learning performance will be best explained by brain area specific pharmacokinetic profiles of receptor densities, drug clearance profiles, or auto-receptor mechanisms that intrinsically downregulate local drug actions (70–72).

In summary, our results provide rare quantitative evidence that a prominent ACh enhancing drug exerts domain specific cognitive improvements of attentional control and cognitive flexibility at a distinct dose range. A major implication of this finding is that for understanding the strength and limitations of pro-cognitive drug compounds it will be essential to test their dose-response efficacy at multiple cognitive domains.

## Acknowledgements

We thank Dr. Carrie K Jones and Jason Russel for helpful feedback throughout the study and about the manuscript. Research reported in this publication was supported by the National Institute of Mental Health of the National Institutes of Health under grants MH123687 (T.W.) The content is solely the responsibility of the authors and does not necessarily represent the official views of the National Institutes of Health.

All authors report no biomedical financial interests or potential conflicts of interest.

## Appendix A Supplementary Information

## SUPPLEMENTAL INFORMATION

## SUPPLEMENTAL METHODS

### Ethics Statement

All animal related experimental procedures were in accordance with the National Institutes of Health Guide for the Care and Use of Laboratory Animals, the Society for Neuroscience Guidelines and Policies, and approved by Vanderbilt University Institutional Animal Care and Use Committee.

### Drug Procedures

For the double blinded drug administration, one experimenter prepared drug doses while another handled injections and observations for potential side effects using a modified Irwin-rating scale. Ratings were assigned on a scale of 0, 1, or 2 per monkey reflecting no change, a slight change or a significant change respectively. Donepezil volumes were separated into vials for storage, and were sonicated and vortexed with sterile saline immediately before injection. Depending on the weight of the animals, the appropriate volume (0.1-0.7 ml) of donepezil was then drawn for the planned injection dose; all daily injections were thus prepared together.

### Visual Stimuli

The behavioral tasks used 3-dimensionally rendered visual objects, so called quaddles, which varied in four visual feature dimensions (shape, color, pattern, and arms of a 3D rendered object) described in detail elsewhere (1). Each visual feature dimension can be parametrically changed which we then used to generate a number of variants, feature values, of each of the mentioned visual features (e.g. up-, and downward bended arms with blunt pointy or flared shape). From here on out, we will refer to the used visual feature spaces as ‘feature dimensions’ and any specific variant of each visual feature as ‘feature values’. During training, all monkeys were exposed to a so-called ‘neutral’ quaddle object composed of a spherical shape, grey color, uniform pattern, and straight arms, which were features values that were never rewarded and served as a null feature value for each dimension. Therefore, to practically achieve objects with only color and pattern feature dimensions, and therefore without shape and arms, we kept the shape and arm dimension constant at the neutral quaddle’s value for shape and arms while having color and pattern feature values that were different from the neutral quaddle’s color and pattern.

### Behavioral Tasks

Monkeys performed a visual search (VS) task and a feature-reward learning (FL) task in each experimental session(1). For each experimental session and for the VS task, we selected randomly 3 feature dimensions from the pool of 4 possible dimensions (shape, color, pattern, arms) and we we chose randomly 3 feature values per dimension (e.g. the 3 shape feature values pyramidal, oblong and cubical) (**Figure 1Bi**).

#### Visual search with different target-distractor similarity

The visual search (VS) task quantified how much visual distractors slow down the detection of a target object and how the distraction varied with the feature similarity of targets and distractors. The task required finding a cued object amongst distractors on the screen by touching it for a minimum of 0.2s. At the beginning of each VS block, the monkey learned which object is the target object in 10 familiarization trials that presented the target object without any distractors. Touching the object triggered fluid reward. The target was always an object that varied in three feature dimensions from the feature values of the neutral object, i.e. a so called 3-D target. This is proceeded by 100 trials, each with a random counterbalanced distribution of 3/6/9/12 distractors. Distractors were also 3-D objects with feature values selected at random and thus could share 0/1/2 features with the rewarded object and could be identical to other distractors within the same trial. If the distractors were dis-similar from the target, independent of the number of distractors, trials may have a pop out effect with the target being easily distinguishable while if distractors were similar to the target, trials may resemble a conjunction search more closely (**Figure 2A**). Objects are presented at random within the intersections of a 4×5 grid (example trials in **Figure 1Bii**).

Individual VS trials are initiated via a 0.3-0.5s long touch to a centrally presented blue square that is 3° radius wide with a sidelength of 3.5 cm (baseline). This was then followed a 0.3-0.5s period where the blue square disappears and there are no objects on screen except for the background image. The task objects are then presented allowing the animal to freely explore for a maximum of 5s (search + selection). During this 5s window, the animal could at any point touch and hold for 0.2s an object in order to select it. The selected object would then prompt both visual and auditory feedback 0.2s after the selection lasting 0.5s. The color of the visual feedback and the pitch of the auditory feedback correspond to the valence of the selected object’s value either signaling a correct or incorrect choice. Correct choices were followed by fluid reward (water) (**Figure 1C**).

The VS task at the beginning and end of the experimental session utilized targets and distractors that were composed of features from the same 3×3 feature space. Targets were never identical between these two blocks but may appear as distractors in the other VS block. Similarly, all distractors were created at random from the same 3×3 feature space as well and therefore would be similar between the two blocks. The background image of the two VS blocks always differed and acted as a cue for the VS rule set but are different in order to prevent the association of the rewarded target object with a particular background image. Thus, the first and second VS search block varied in the target object pulled from the same 3×3 feature space, the background image, as well as the timing of their occurrence being at the start or the end of the daily session (**Figure 1Aii**).

#### Feature-reward learning at different distractor load

The feature-reward learning (FL) task quantifies how fast and accurate subjects adjust to changing reward rules, indexing cognitive flexibility. The task required monkey to learn by trial-and-error which object feature is associated with reward. The same feature remained rewarded within blocks of 35-60 trials. Monkeys had to choose one amongst three objects (1 target and 2 distractors) where a single feature value in a single feature dimension is linked to reward with a p = 0.85 reward probability. Distractors contain the same dimensions as the target but have different, non-repeated feature values. All objects are presented in 1 of 8 possible locations randomly, all with 17 degrees eccentricity from the central touch location (example trials in **Figure 1Bii**). With one experimental session we ran 21 FL blocks. The feature-reward association must be learned through trial and error and may switch after 35, 40, or 45 trials from the start of the block if the learning criterion is reached (80% over 10 trials) or in 60 trials otherwise (uniform max FL block trial number). Block changes are un-cued but can be inferred if there is a change in the object feature dimensions presented and the newly rewarded feature value may be in the same dimension or a different dimension relative to the previously rewarded feature value; the two types of shifts are semi-randomly determined to occur in similar frequencies. The temporal structure and sequence of epochs in the FL task is the same as the VS task.

The structure of the trials within the FL task was very similar to that of the VS task. Trials are initiated in a similar manner via a 0.3-0.5s touch on a central blue square. This is followed by a 0.3-0.5s period where the blue square is not present and task objects have not yet been made visible yet. Three task objects are then presented for up to 5 sec during which at any point the subject is allowed to make a 0.2s touch and hold on an object to select it. Following a 0.2s delay after the selection of an object, auditory and visual feedback as well as potentially fluid reward are presented for 0.5 s. The pitch of the auditory feedback and the color of the visual feedback vary depending on the presence of reward and not on making a high reward probability choice (**Figure 1C**).

### Neurochemical Quantification of Drug Effect

To confirm the bioactivity of donepezil in the brain we measured the neurochemistry in the prefrontal cortex and the head of the caudate nucleus after IM administering a low (0.06 mg/kg) and high (0.3 mg/kg) dose of donepezil. We used microprobes that sampled the local neurochemical milieu with the principles of solid phase micro-extraction (SPME) probes (2) previously shown to provide comparable and complimentary outcomes to micro-dialysis (3,4). These probes sample the drug and metabolites of the neurotransmitters (e.g choline) via diffusion until an equilibrium is reached with the extracellular concentrations. The detailed procedures used here are described in (5). In brief, for each brain area a microdrive was prepared holding a cannula and SPME probe inside it, as well as a microdrive with an electrode to record activity prior to SPME sampling. The electrode was driven to the target location in prefrontal cortex / striatum. The target location was confirmed by measuring spiking activity of neurons from the electrode. The cannula shielded SPME was then lowered to just above the target area and the SPME probe was then exposed to gray matter of the target area for 20 minutes before being retracted into their respective cannula and drive back out of the brain. Samples were then stored in a -80°C freezer, stored for less than 2 weeks and shipped overnight in dry ice to Waterloo, Ontario (Canada) where they were desorbed and underwent liquid chromatography separation and mass spectrometry quantification.

The SPME probes were desorbed into 50 µL of acetonitrile/methanol/water 40:30:30 solution containing 0.1% formic acid and internal standard citalopram-D_6_ at 20 ng/mL for 1 h with agitation at 1500 rpm. The LC-MS/MS analysis was carried out using an Ultimate 3000RS HPLC system coupled to a TSQ Quantiva triple quadrupole mass spectrometer (Thermo Fisher Scientific, San Jose, CA, USA). Data acquisition and processing were performed using Xcalibur 4.0 and Trace Finder 3.3 software (Thermo Fisher Scientific, San Jose, CA, USA). The chromatographic separation employed Hypersil Gold C18 column, 50 × 2.1 mm, 1.9 μm particle size (Thermo Scientific, Ashville, NC, United States) held at 35°C. The aqueous mobile phase (A) consisted of water/acetonitrile/methanol 90:5:5 with 0.1 % formic acid, while the organic mobile phase (B) consisted of acetonitrile/water 90:10 with 0.1% formic acid. The following chromatographic gradient at a flow rate of 400 µL/min was applied (%B): 0-0.5 min 0 %; 0.5-3 min linear gradient to 100 %; 3-3.65 min held at 100 %; 3.65-3.7 min linear gradient to 0 %; re-equilibration at 0 % until 4.5 min. The injection volume was 5 μL. The MS/MS analysis was performed in positive ionization mode under selected reaction monitoring (SRM) conditions; for the analyte donepezil the quantifier transition monitored was m/z 380.3 -> 243.2 and the qualifier transition was 380.3 - > 91.1, while one transition at m/z 331.1 -> 109.1 was monitored for internal standard citalopram-D_6_. The capillary voltage was set at 3.5 kV, with the remaining electrospray source conditions set to the following values: vaporizer temperature 358 °C, ion transfer tube temperature 342 °C, sheath gas pressure 45, auxiliary gas pressure 13, and sweep gas pressure 1 (arbitrary units). The instrumental stability throughout the sequence was monitored by analysis of an instrumental QC sample consisting of the target analyte and internal standards spiked into a neat desorption solvent at 20 ng/mL.

The concentration of donepezil in brain tissue was determined using a modified external surrogate matrix-matched calibration approach developed in previous work (5–7). The surrogate matrix consisted of agarose gel (1% agarose in PBS solution, *w/v*) mixed with lamb brain homogenate in the ratio 1:1 (*v/w)*. Prior to combining the agarose gel with the brain homogenate, the latter was spiked with donepezil in the concentration range 5-750 ng/g. Extractions were carried out in static mode from 1g of the matrix with extraction time matching the *in vivo* experiments. The probes were subsequently rinsed with water and desorbed into 50 µL of the desorption solvent containing internal standard citalopram-D_6_ at 20 ng/mL. The analytical response in the form of relative peak area ratios (analyte to internal standard) was converted to amounts extracted by employing an instrumental calibration curve consisting of donepezil in neat desorption solvent in the range 0.1-100 ng/mL. The resulting matrix-matched calibration curve was expressed as amounts extracted in the function of concentration in tissue. A weighted linear regression equation was fitted to the analytical response in the function of concentration. Limits of quantitation (LOQ) were determined as the lowest concentration of analyte producing a signal to noise ratio ≥ 5, with a relative standard deviation (RSD) of 4 replicate measurements below 20%, and accuracy within 20%. Accuracy was calculated as the relative percent error of concentrations of analytes in the calibrator samples determined experimentally with the use of calibration curves versus theoretical (spiked) concentrations (8).

A single, fourth, *Macaca mulatta* (male, 8 years old) with an implanted recording chamber above the left hemisphere was chaired, head-fixed and performed the VS task (data not included in analysis) to emulate performance by the other 3 subjects. Details about the surgical implantation of the recording chamber and headpost are reported in (5). Performance of the VS, virtually identical to the VS task reported above, was done with eye saccades using a Tobii Spectrum eye tracker instead of touch screen. Subject underwent 6 instances of both 0.06 and 0.3 mg/kg donepezil dosing in the primate chair at the same time of day as the other 3 NHPs received donepezil. Injections were done after the animal had been performing one VS block for roughly 20min, followed by a 15min period of quiet wakefulness after which they proceeded to do a second VS block. SPME sampling events took place once at the beginning of each VS block with probes being exposed to tissue for 20min in both instances.

During each SPME sampling event heart rate (HR) was monitored using a pulse oximeter (PalmSAT 2500, Nonin Inc, MN), with the sensor clipped at the ear lobe of the subject and a sampling rate of 0.25 Hz. HR data was collected 20 min before task start both before and after donepezil injection. The data was smoothed with an centered 8 sample window (40 sec) with 1 sample shifts (4 sec) and normalized to the average HR 5 min before task start.

### Literature Survey

In order to place our results within the broader published work, we identified 9 papers involving donepezil and non-human primates (9–17). These papers have relevant details such as the task(s) performed, donepezil dosage and administration method and more extracted, where appropriate (**Table S1**). Notably, 6 of the identified papers provided donepezil in conjunction with other pharmaceutical agents such as Scopolamine. The papers were found by conducting an online search of the NIH (PubMed) database, as well as google scholar. The keyword search terms of ‘Donepezil’, ‘Aricept’ and ‘E2020’ were used with the terms ‘NHP’, ‘monkey’, ‘primate’, ‘cognition’, and ‘brain’ or some combination of them. We did not consider 8 studies that utilized donepezil in primates as they lacked a cognitive component. They did however provide insight in dosing ranges for different dosing routes, dose-limiting side effects and donepezil’s kinetics (18–25).

### Data Analysis

All behavioral analysis was completed using MATLAB (Mathworks Inc., MA).

#### Analysis of Visual Search

The set size effect of the VS performance was either defined as proportion of correct trials by the distractor number or by the average t-d similarity of trials. The set-size effect was estimated by a linear regression which is specified as either utilizing distractor number or t-d similarity. The average t-d similarity of a trial was calculated by averaging the number of shared feature values (0/1/2) of all distractors in said trial to the target. Reaction times, referred to as choice RTs for the VS task, were defined as the time from the initiation of a trial by pressing and holding the central blue square to the initiation of touch to the selected object leading to feedback. Reaction time data only takes into consideration rewarded trials. Descriptive statistics are provided as means with ±SEMean unless specified otherwise. Similarly, error bars in figures are either mean ±SEMean or median ±SEMedian unless specified otherwise. After pooling data from all three subjects, the measure of interest is averaged across appropriate trials or blocks to get a per session value.

#### Analysis of Feature-Reward Learning

FL blocks were either labeled as ‘low distractor load’ if no distracting feature dimension was present, or as ‘high distractor load’ if a single distracting feature dimension was presented alongside the feature dimension to which the rewarded feature value belonged to. We calculated learning curves by averaging smoothed trial-wise performance aligned to block reversals. We defined learning speed by calculating at which trial, since block start, the subjects started performing at ≥80% over 10 trials, the maximally rewarded object. This trial was termed the ‘learning point’ (LP). For analysis, blocks were excluded if the monkey took a break of at least several minutes. Furthermore, blocks were excluded where the LP was calculated to be trial 1 (reflecting ≥80% performance in the first 10 trials since reversal) as well as if the LP occurred beyond the 40^th^ trial. Reaction times, referred to as choice RTs for the FL task, were temporally defined the same as for the VS task and also only include rewarded trials.

Perseverative errors were defined as two or more consecutive choices of low probability rewarded objects with at least 1 shared feature value. Analysis of perseverative errors for feature values in the same feature dimension as the target feature are separated from those where the perseverated feature value was in the distracting dimension. For perseverative errors to occur in the distractor dimension, the block is required to contain a distracting dimension to begin with and is therefore necessarily a high distractor load block. Perseverative errors in the target feature dimension could occur in both low and high distractor load blocks.

### Statistical Analysis of Drug Effects

Comparisons between vehicle and donepezil doses (0.06, 0.1 and 0.3 mg/kg) were done for all doses combined followed by post-hoc pair-wise statistics with multiple comparisons corrections unless specified otherwise. Probability level of less than 5% (p < 0.05) was considered statistically significant.

## SUPPLEMENTAL RESULTS

### Overall Visual Search Performance

We performed and report here the results of various analysis to evaluate the overall performance of the animals on the tasks, or to test specific performance metrics that provide a more comprehensive overview of how the drug conditions did or did not affect task performance.

For the visual search task, 10 familiarization trials with no distractors were presented prior to each of the two visual search blocks. The reaction times to detect these single objects on the screen will be referred to as speed of processing (SoP). They were completed in 628 ms ±133 (Ig: 616 ms ±8.5; Wo: 693 ms ±6.6; Si: 588 ms ±5.3) with the first block having faster SoP at 611 ms ±7.7 relative to the second block with 646 ms ±4.8 (p < .001)(**Figure S1A**).

On average, monkeys performed the VS task with 84.4% (± 0.54) accuracy (Monkey Ig: 85.2% ±0.81; Wo: 88.3% ±0.94; Si: 79.8% ±0.97) and with 1158 ms ±9.7 search times (Ig: 1281 ms ±18.3; Wo: 1171 ms ±15.9; Si: 1020 ms ±13.3). Increasing numbers of distractors slowed search RTs, with 3/6/9/12 distractors having 1019 ms, 1216 ms, 1409 ms, and 1552 ms search times respectively (Distractor Number F(3,1240) = 241.32, p < .001) as well as decreasing accuracy, with 3/6/9/12 distractors having 91.7% ±0.6, 87.1% ±0.6, 82.9% ±0.8, and 80.0% ± 0.9 accuracy respectively (all pair-wise comparisons were significant using Tukey’s HSD multiple comparisons test among proportions at an alpha of 0.05, except for 0.1 mg/kg donepezil dose and vehicle). Search RTs did not vary significantly with regards to VS block number (Block Number F(1,1240) = 3.18, n.s.) (**Figure S1B**) while trial outcomes did vary significantly with VS block number (*X*^2^(1, N_1_ = 14500, N_2_ = 16700) = 40.0, p < .001) (**Figure 1D**). This difference in performance between the two VS blocks may be due to fatigue, as reflected by the significantly reduced SoP, or otherwise satiation.

Both the change in search time and performance by distractor number were fit by a linear regression, revealing that each additional distractor increased search duration on average by 60 ± 1.6 ms (Ig: 72 ± 2.7 ms/distractor; Wo: 57 ± 2.8 ms/distractor; Si: 49 ± 2.1 ms/distractor) as well as decreasing performance by 1.3% ± 0.1 per additional distractor (Ig: 1.2% ± 0.1; Wo: 0.9% ± 0.1; Si: 1.8% ± 0.1) (inlets in **Figures 1D and S1B** show individual monkey fits for vehicle). The set size effect on search RT was on average larger in the first than the second VS block (first VS block: 63 ms/distractor; second VS block: 56 ms/distractor; p = .0254; Ig: p = .0604; Wo: p = .0401; Si: p = .7199). The set size effect on performance was on average the same in the first and the second VS block (first VS block: -1.3% change in performance per distractor; second VS block: -1.3% change in performance per distractor; p = n.s.; Ig: p = n.s.; Wo: p = n.s.; Si: p = n.s.).

We analyzed how the similarity of distractors with the target influenced search RT and performance. Distracting stimuli could have 0, 1 or 2 shared feature values with the target and the thus some trials could provide a greater challenge for the monkeys given the average target-distractor similarity (t-d similarity)(**Figure 2A**). We found that search RT increased with average t-d similarity from 1227 ms ±9 to 1410 ms ±7 and 1334 ms ±17 for low, medium and high t-d similarity respectively (Similarity F(2,14467) = 107.1, p < .001)(**Figure S4A**). VS performance decreased with t-d similarity from 92.9% ±0.4 to 85.5% ±0.3 and 81.6% ±1.0 for low, medium and high t-d similarity respectively (Similarity F(2,672) = 16.17, p < .001) (**Figure 2B**). Both distractor number, and t-d similarity impact VS performance significantly (Distractor number F(3,16688) = 50.25, p < .001; T-D similarity main effect p < 0.001)(**Figure 4B-C**) but no significant interaction was found between the two variables (T-D Similarity x Distractor Number F(6,16688) = 1.16, n.s.)(**Figure 4D**) with VS RT showing a similar relationship with significant main effects (distractor number main effect p < 0.001; T-D similarity F(2,16688) = 55.24, p < .001) but not interaction (F(6,16688) = 1.16, n.s.)(**Figure 2D**). Individual sessions also showed no strong correlation between the set size effect of performance by distractor number relative to the set size effect of performance by t-d similarity (Pearson, n.s.)(**Figure S3**).

### Overall Feature Reward Learning Performance

For the feature reward learning (FL) task, monkeys reached learning criterion on average in 63 ±1% of the 21 daily learning blocks (Ig: 71 ±1%; Wo: 61 ±2%; Si: 56 ±1%) once exclusion criteria were applied (see methods). Learning criterion was reached more often in the low distractor load than high distractor load blocks with proportion of learned blocks being 70% and 56% respectively (Ig: 80 vs 62% of blocks; Wo: 66 vs 56%; Si: 63 vs 49%). Average learning curves for low and high distractor load blocks of each individual monkey, as well as the average across monkeys is provided in **Figure 3A**. Monkeys reached the learning criterion on average within 12.5±0.2 and 15.6±0.2 trials in the low and high distractor load condition (Ig: 9.9±0.2 and 14.9±0.3; Wo: 13.5±0.4 and 17.0±0.4; Si: 14.9±0.4 and 15.0±0.4). The average choice reaction time of a correct FL trial was 986 ±2 ms with faster reaction times in the low than high distractor load blocks (p < .001; 965 ±3 ms and 1013 ±3 ms respectively).

### Visual search performance with donepezil

The speed of processing (SoP; reaction time to a single object during familiarization trials) showed a significant main effect of block number (Block Number F(1,424) = 6.29, p < .001), as well as a significant main effect of drug condition (Condition F(3,424) = 15.16, p < .001)(**Figure 2B**). Pair-wise statistics comparing the first block SoPs of the control condition and 0.06, 0.1 and 0.3 mg/kg doses (Tukey’s, n.s, n.s, and p < .001 respectively) suggests that the main effect of condition is driven by the 0.3 mg/kg dose SoP.

### Feature learning performance with donepezil

For the low distractor load condition the proportion of learned blocks were on average 72.3% ±1.4, 75.2% ±2.4, 78.0% ±3.9 and 75.9% ±3.2 in the vehicle, and 0.06 / 0.1 / 0.3 mg/kg) days, which was not significant (n.s.). Similarly, for the high distractor load condition the proportion of learned blocks was 60.0% ±1.5, 74% ±4.0, 62.3% ±3.8 and 65.0% ±4.2 in the vehicle, and 0.06 / 0.1 / 0.3 mg/kg) days (n.s.). Comparisons between the proportion of blocks learned in low and high distractor load conditions revealed a significant difference for vehicle and drug conditions with more blocks learned in the low distractor load condition than in the high distractor load condition (*X*^2^ values contrast 1D vs 2D learning blocks for vehicle, 0.06, 0.1, and 0.3 mg/kg conditions: *X*^2^(1, N_1_ = 1757, N_2_ = 1750) = 58.6, p < .001; *X*^2^(1, N_1_ = 222, N_2_ = 219) = 8.2, p = .004; *X*^2^(1, N_1_ = 219, N_2_ = 222) = 12.6, p < .001; *X*^2^(1, N_1_ = 209, N_2_ = 190) = 5.6, p = .018 for vehicle, 0.06, 0.1 and 0.3 mg/kg doses respectively). Monkey Ig had a higher overall proportion of blocks learned than both Monkey Wo and Si with vehicle (p < .001), however, there were no statistically significant differences between monkeys between low and high distractor loads in vehicle or any drug conditions (n.s.).

In addition to learning speed we also analyzed in detail the choice reaction times across drug conditions. Relative to the low distractor load condition, in the high distractor load condition, FL choice RTs slowed from 993 ±11 to 1060 ±14, from 964 ± 31 to 1048 ±33, from 988 ±27 to 1015 ±36, and from 1126 ±29 to 1167 ±31 ms for the vehicle, 0.06, 0.1 and 0.3 mg/kg donepezil doses respectively (**Figure 3E**).

There was also significant inter-subject variability in choice RTs with monkey Si having significant faster choice RTs in the FL task (Subject F(2,1052) = 183.53, p < .001)(**Figure S2C**) as well as a significant monkey-drug interaction (F(6,1052) = 3.5, p = .002). Alongside the general slowing with the 0.3 mg/kg donepezil dose (see main text), we found in a pair-wise analysis a significant slowing of search RTs with the 0.3 mg/kg donepezil dose for monkey Si (Tukey’s, p < .001), and a significant faster search RTs with the 0.1 mg/kg donepezil dose for monkey Wo (Tukey’s, p = .003).

The main text reports the length of consecutive, perseverative errors. Perseverative errors may occur in the same dimension as the target feature value (12% of all errors), possible in low and high attentional load blocks, or they may occur in the distracting dimension (26% of all errors) only possible in high attentional load blocks. The proportions of perseverative errors within the target dimension were 12% ±1, 11% ±2, 12% ±2 and 11% ±2 for vehicle, 0.06, 0.1 and 0.3 mg/kg donepezil doses (n.s.), while within the distracting dimension they were 26% ±0, 23% ±6, 22% ±2, and 24% ±2 for vehicle, 0.06, 0.1 and 0.3 mg/kg donepezil doses (n.s.).

We next analyzed whether donepezil modified how flexible subjects learned a new target feature depending on whether the target feature was from a novel feature dimension and whether the target was a previous distractor. First, we asked whether donepezil modulates learning differently depending on whether a newly rewarded (target) feature values belonged to the same feature dimension as the target feature in the previous block, or to a new target feature dimension. This analysis quantifies whether learning a new feature set was easier or more difficult than re-assigning a reward association within the previously relevant feature set. In our task a shift to a new target feature of a new dimension should be easier because it occurred by presenting new objects that were not shown in the previous block. We thus compared learning speed for blocks where the rewarded feature dimension was not presented in the previous block and blocks where the rewarded feature value was from the same dimension as the previously rewarded feature. We found that donepezil did not alter learning for block transitions to ‘new target feature dimensions’ versus ‘another feature of the same dimension’ (Condition F(3,2708) = 0.55, n.s.; Block Switch F(1,2708) = 2.7, n.s.; Condition x Block Switch F(3,2708) = 1.15, n.s.).

Secondly, we quantified whether donepezil modulated how subjects learned a new target feature value when that target feature was a distractor in the previous block. Difficulties in attending a previous distractor is sometimes referred to as latent inhibition. There were only few learning blocks available in which the target feature dimension was a distracting feature dimension in the previous block which we contrasted to blocks where the rewarded feature dimension was not presented in the previous block. We found that donepezil did not alter learning speed for blocks where the ‘target was a previous distractor’, versus when the ‘target was a new feature’ (Condition F(3,1450) = 0.31, n.s.; Block Switch F(1,1450) = 0.02, n.s.; Condition x Block Switch F(3,1450) = 0.2, n.s.).

## SUPPLEMENTAL TABLES

**Table S1.** A summary of the literature testing donepezil’s cognitive effects in nonhuman primates.

| Relevant Task(s) | Subject Details | Dosage & Administration | Cognitive Domain | Relevant Results | Reference |
| --- | --- | --- | --- | --- | --- |
| Object retrieval detour (ORD) cognition test | Macaca mulatta (male and female) | 0.3, 0.56, 1, 1.8, 3, 5 mg/kg PO* | Reasoning & problem solving (exec function) | Significant interaction of trial type (easy vs difficult) and treatment. Main effect of treatment on the difficult condition | Vardigan et al., 2015 (9) |
| Paired-associates learning (PAL); Continuous-performance task (CPT) | Macaca mulatta (18 males) | 0.3-3 mg/kg PO (PAL task); 0.1-0.25 mg/kg IM (CPT task)* | PAL: Working memory; CPT: Attention/vigilance (exec functioning) | Attenuated scopolamine-induced impairments in PAL (at 1.0 and 3.0 mg/kg PO) and CPT (at 0.25 mg/kg IM) | Lange et al., 2015 (10) |
| Delayed matching-to-sample (DMTS) task | Macaca mulatta (4 aged male, 3 aged female) | 0.003-0.2 mg/kg PO* | Working memory | Enhanced DMTS accuracy in 'long delay' condition at 0.01, 0.025, 0.05, 0.1 and 0.2 mg/kg doses. No changes in ITI or choice latency | Callahan et al., 2013 (11) |
| Self-ordered spatial search | Macaca fascicularis (6 females; ~15 years old) | 3 mg/kg PO* | Working memory, Attention/vigilance | Attenuated the scopolamine-induced impairments in the self-ordered spatial search task | Uslaner et al., 2013 (12) |
| Delayed matching-to-sample (DMTS) task | Macaca mulatta (17 male and 16 female; average ~18 years old) | 10, 25, 50, 100 ug/kg IM<br>50-400 ug/kg PO* | Working memory | Accuracy in long delay condition with IM administration was improved | Buccafusco et al., 2008 (13) |
| Delayed matching-to-sample (DMTS) task | Macaca mulatta (8 male & 9 female; 9-29 years old) | 10, 25, 50, 100 ug/kg IM | Working memory | Accuracy increased in medium and long delayed trials (with 25 ug/kg dose being the most efficacious) | Buccafusco & Terry 2004 (14) |
| Oculomotor delay response task (ODR); Visually guided saccade task (VGS) | Macaca mulatta (male; 5 ~5 years old and 5 ~20 years old) | 50, 250 ug/kg IV | ODR: Working memory; VGS: attention/vigilance | ODR performance was improved in aged monkeys (not young monkeys). No changes reported in VGS (assay of motor performance, not cognition) | Tsukada et al., 2004 (15) |
| Delayed matching-to-sample (DMTS) task | Macaca mulatta (male & female; >20 years old) | 0.01, 0.025, 0.05, 0.1 mg/kg IM | Working memory | Accuracy improved independent of drug dose but dependent on delays (improvement occurred in medium and long delays) | Buccafusco et al., 2003 (16) |
| Spatial delayed response task (SDRT); Visual recognition task (VRT) | Macaca mulatta (9 males) | 0.01-1.75 mg/kg IM (SDRT task); 0.003-0.06 mg/kg IM (VRT task)* | SDRT: Working memory; VRT: attention/vigilance | SDRT: Donepezil rescued effects of scopolamine in difficult trials<br>VRT: Performance was enhanced with donepezil pre-treatment (by ~10%) | Rupniak et al., 1997 (17) |

\*: Other drugs co-administered at some/all doses.

**Table S2.** A summary of observed dose-limiting side effects. The effects of donepezil (0.06, 0.1 and 0.3 mg/kg IM) on autonomic and somatomotor system function were evaluated. The mean score of 3 monkeys was classified as follows: - no effect; + 0-0.15; ++ 0.16-0.3; +++ 0.31-0.45.

| Donepezil |  | 0.06 mg/kg |  | 0.1 mg/kg |  | 0.3 mg/kg |  |
| --- | --- | --- | --- | --- | --- | --- | --- |
|  | Observation | Pre-task | Post-task | Pre-task | Post-task | Pre-task | Post-task |
| Autonomic Nervous System | Salivation | - | - | - | - | - | - |
|  | Lacrimation | - | - | - | - | - | - |
|  | Urination | - | - | - | - | - | - |
|  | Defecation (amount) | - | - | - | - | + | - |
|  | Defecation (consistency) | - | - | - | - | + | - |
|  | Emesis | - | - | - | - | - | - |
|  | Miosis | - | - | - | - | - | - |
|  | Mydriasis | - | - | - | - | - | - |
|  | Ptosis | - | - | - | - | + | - |
|  | Exophthalmos | - | - | - | - | - | - |
|  | Piloerection | - | - | - | - | - | - |
|  | Respiratory Rate | - | - | - | - | - | - |
|  | Yawn | - | - | - | - | + | - |
|  | Vasodilation | - | - | - | - | - | - |
|  | Vasoconstriction | - | - | - | - | +++ | - |
|  | Irritability | - | - | - | - | - | - |
|  | Body Temp. | - | - | - | - | - | - |
| Somatomotor Systems | Physical Appearance | - | - | - | - | +++ | - |
|  | Tremor | - | - | - | - | - | - |
|  | Leg Weakness | - | - | - | - | - | - |
|  | Catalepsy | - | - | - | - | - | - |
|  | Visuo-Motor Coordination | - | - | - | - | - | - |
|  | Posture | - | - | - | - | +++ | - |
|  | Unrest | - | - | - | - | - | - |
|  | Stereotypies | - | - | - | - | - | - |
|  | Arousal | - | - | - | - | - | - |
|  | Sedation | - | - | - | - | +++ | - |
|  | Oral Dyskinesia | - | - | - | - | ++ | - |
|  | Bradykinesia | - | - | - | - | + | - |
|  | Dystonia | - | - | - | - | - | - |

## SUPPLEMENTAL FIGURES

**Figure S1.**
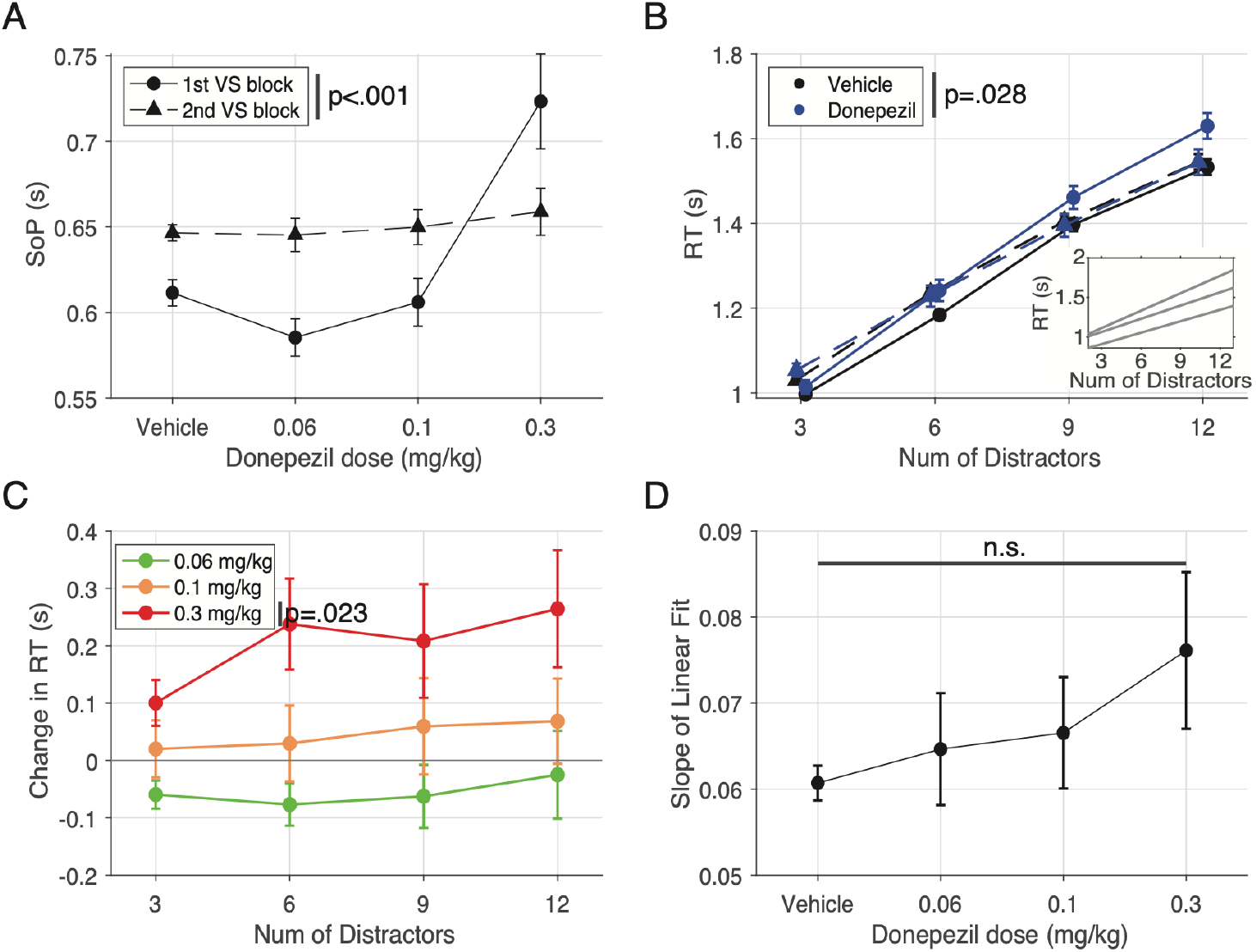
Search reaction time in the visual search task and its relationship with distractor number. **A**. The average speed of processing (SoP), for each condition separated by block. The SoP is significantly changed between the first and second VS block (Block Number F(1,424) = 6.29, p < .001) as well as between conditions (Condition F(3,424) = 15.16, p < .001; ANOVA). The average SoP in the first VS block is significantly slowed with a 0.3 mg/kg dose of donepezil (Tukey’s, p < 0.001). **B**. Average search RT per distractor number for vehicle and all donepezil doses combined, both separated by the first vs second VS block. The number of distractors slowed search RTs (Distractor F(3,1722) = 333.1, p < .001) while the VS block number did not significantly impact search RTs (Block F(1,1722) = 0.64, n.s.). Donepezil administration, averaged over all doses, had a significant effect on search RT (Condition F(1,1722) = 4.83, p = .028), in particular in the first VS block. The inlet shows individual monkey average search RT linear fits. **C**. The difference in search RT by distractor number between donepezil 0.06, 0.1, 0.3 mg/kg doses and vehicle for the first VS block (F(3,896) = 15.15, p < 0.001) with the 0.3 mg/kg dose having significantly slower search RT than vehicle (p = .023). Error bars are standard deviations. **D**. The set size effect of search RT by distractor number for each condition. No significant difference was observed between drug conditions and vehicle.

**Figure S2.**
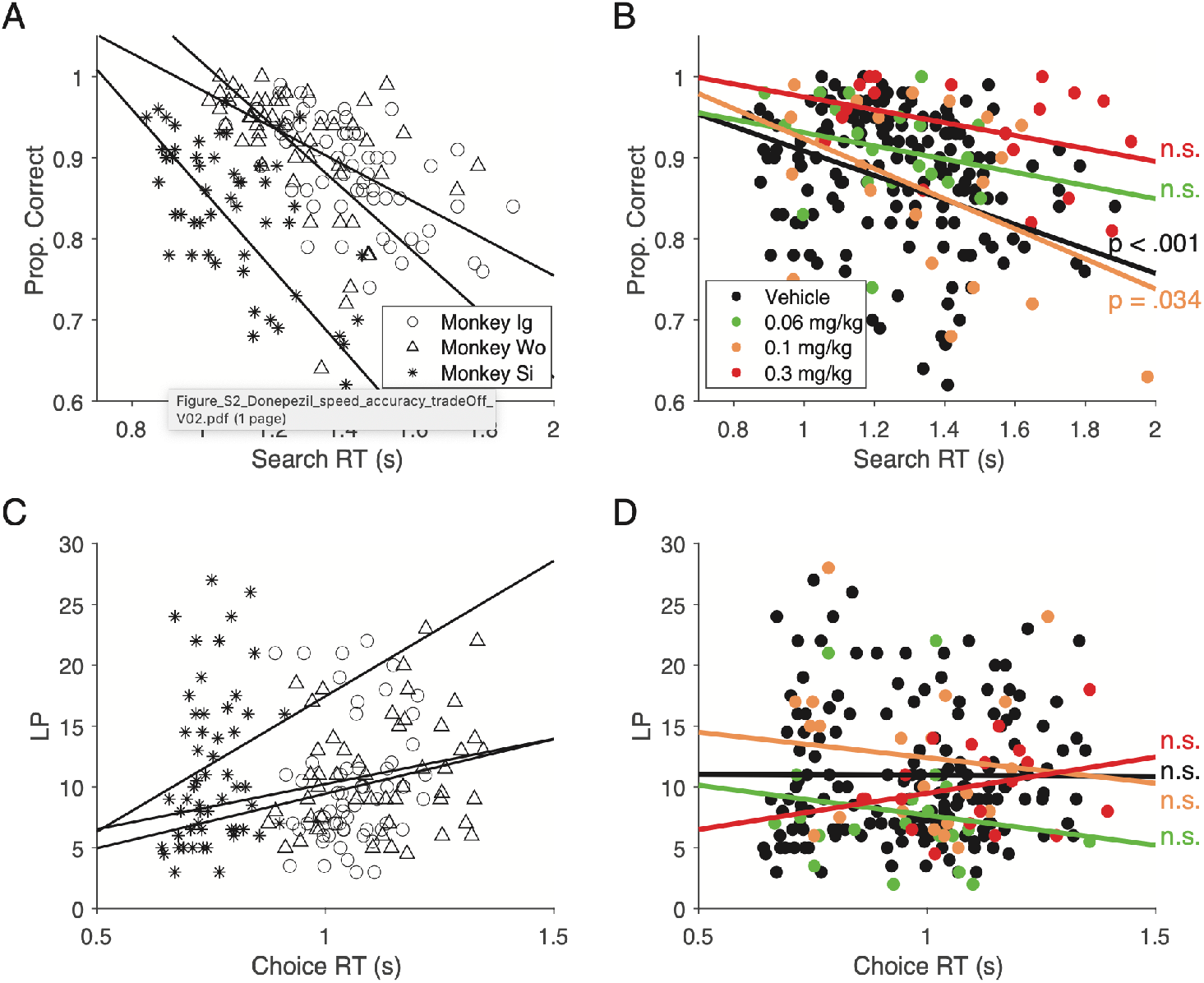
The relationship between performance and reaction times in both the visual search task and feature learning task. **A**. The session-wise correlation between VS performance and search RT by individual monkey. No difference between monkeys was found. **B**. Same as A but for all monkeys combined and separated by condition. Only vehicle and 0.1 mg/kg doses had a significant correlation, however no significant change in correlation relative to vehicle was found. **C**. Similar to A but looking at the correlation between FL performance (learning speed) and choice RT. Monkey Si was found to have significantly faster choice RTs (Subject F(2,1052) = 183.53, p < .001). **D**. The same as C but for all monkeys combined and separated by condition. No conditions exhibited significant correlations.

**Figure S3.**
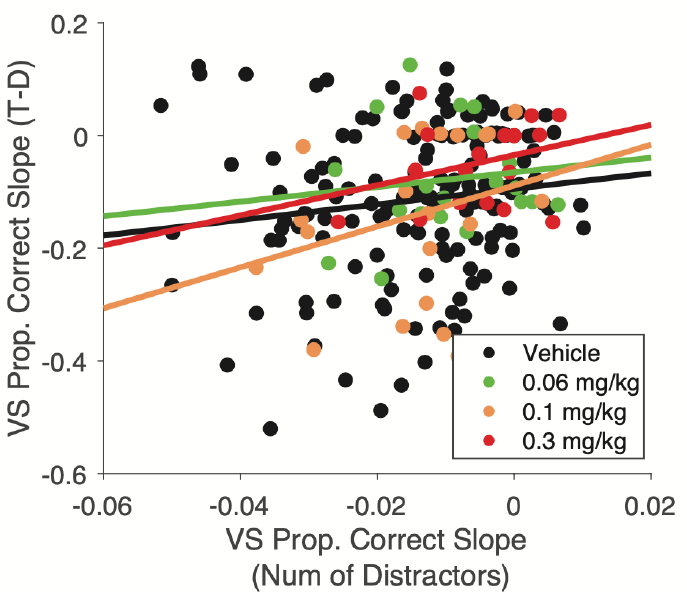
The relationship between the set-size effect of visual search performance as a function of distractor number versus target-distractor similarity. Session-wise linear fits to performance by distractor number (*x-axis*) and target-distractor similarity (*y-axis*). There was no significant correlation at any condition.

**Figure S4.**
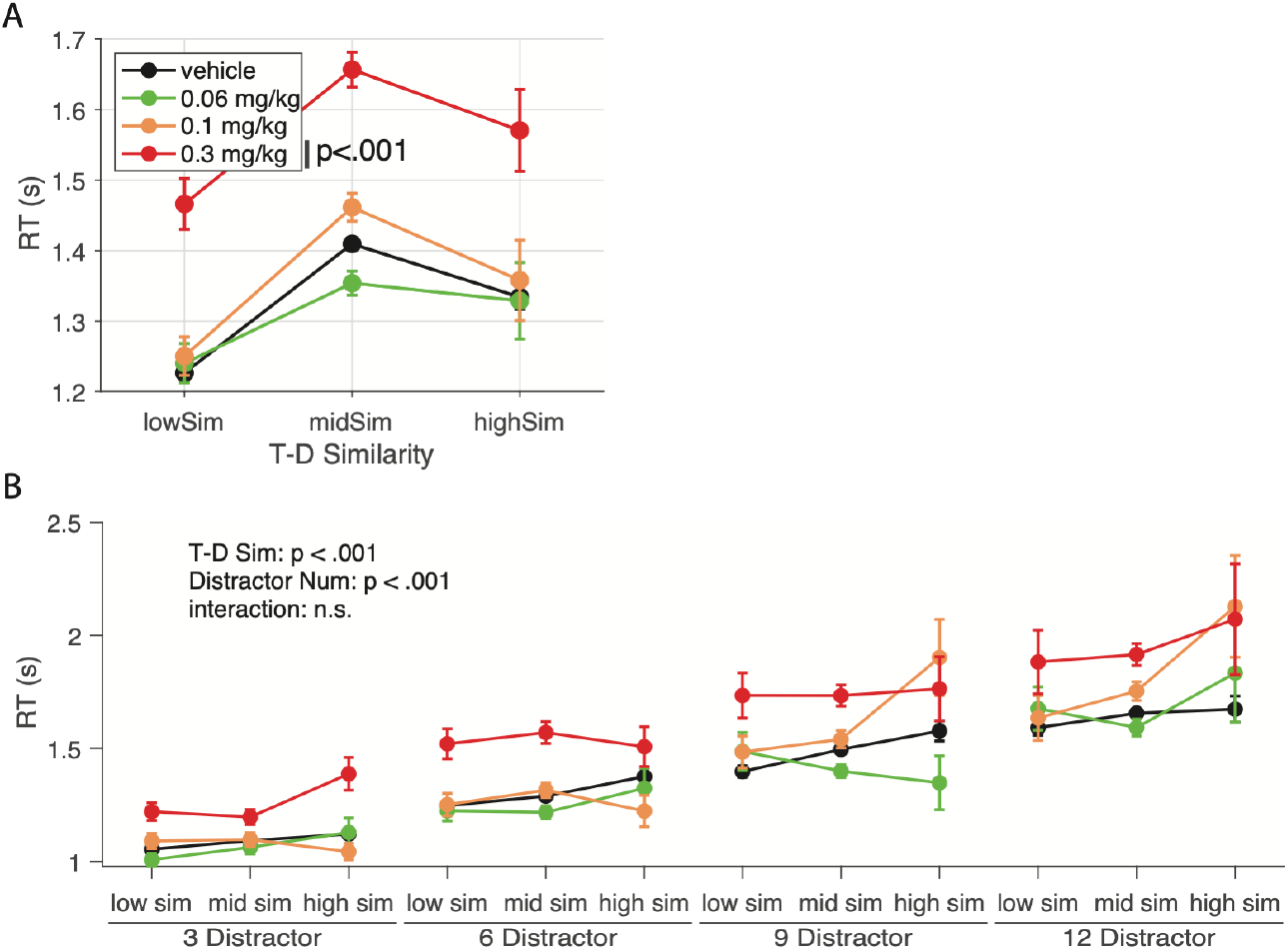
Search reaction times within the visual search task as a function of target-distractor similarity and distractor number. **A**. Search reaction time plots as a function of t-d similarity instead of distractor number (as was the case in **Figure S1**) for vehicle and all donepezil doses. There was a significant main effect of condition with the 0.3 mg/kg donepezil dose being significantly different from vehicle (F(3,267) = 7.75, p < .001; Tukey’s, p < .001). **B**. Visualization of the combined effect of distractor number and t-d similarity on search RT. From left to right, each cluster of lines represents increasing distractor numbers while data within each line represents low, medium and high t-d similarity from left to right respectively. Both distractor number (F(3, 14458) = 294.93, p < .001) and t-d similarity (F(2,14458) = 16.8<u>7</u>, p < .001) impact VS performance with no significant interaction (F(6, 14458) = 1.19, n.s.).

